# Extracellular calcium modulates pollen tube growth and guidance in *Arabidopsis thaliana*

**DOI:** 10.64898/2026.02.07.704530

**Authors:** Kumi Matsuura-Tokita, Yoko Mizuta, Daisuke Kurihara, Tetsuya Higashiyama

**Affiliations:** Department of Biological Sciences, Graduate School of Science, The University of Tokyo, Hongo, Bunkyo-ku, Tokyo 113-0033, Japan; Institute of Transformative Bio-Molecules (WPI-ITbM), Nagoya University, Furo-cho, Chikusa-ku, Nagoya, Aichi 464-8601, Japan; Institute for Advanced Research (IAR), Nagoya University, Furo-cho, Chikusa-ku, Nagoya, Aichi 464-8601, Japan

**Keywords:** calcium imaging, GCaMP6, pollen tube, vegetative nucleus, live imaging, guidance

## Abstract

In angiosperms, pollen tubes deliver sperm cells to the ovule and communicate with the external environment as they elongate through the pistils. Although pollination alters Ca^2+^ conditions within the pistil, the effects of extracellular Ca^2+^ fluctuations on pollen tube growth and guidance remain largely unknown. In this study, we visualized intracellular Ca^2+^ dynamics using a semi-*in vivo* assay with the Ca^2+^-sensitive fluorescent protein GCaMP6s to investigate how pollen tubes respond to changes in extracellular Ca^2+^ levels. We found that the Ca^2+^ levels in the apical region of the pollen tubes reflected the extracellular Ca^2+^ concentrations. The pollen tube growth rate increased depending on the Ca^2+^ concentration in the growth medium. However, excessive Ca^2+^ affected the polar growth of pollen tubes. At elevated Ca^2+^ concentrations of 10 mM, the pollen tube exhibited coiling behavior and failed to maintain directional growth toward the ovule. Moreover, we provided the first evidence that Ca^2+^ oscillations are not restricted to the apical region but propagate as a wave, reaching 30–50 μm from the apex toward the basal regions. As the pollen tube approached the ovule, it coincided with a substantial elevation in Ca^2+^ levels, which appeared to drive the accelerated nuclear migration toward the tube apex. Our findings demonstrate that the extracellular Ca^2+^ environment directly regulates intracellular Ca^2+^ levels in pollen tubes, thereby influencing their growth and guidance.

## Introduction

Ca^2+^ plays a critical role in cellular growth. In animals, extracellular Ca^2+^ influx and intracellular Ca^2+^ gradients regulate both the elongation and directional growth of neuronal axons [1][2]. In plant cells, extracellular Ca^2+^ modulates intracellular Ca^2+^ concentration and the polarized growth of fungal hyphae and root hair cells [3][4]. A similar Ca^2+^ gradient has also been observed in pollen tubes during plant reproduction. In angiosperms, pollen grains act as male gametophytes that contain sperm cells. Upon pollination on the stigma, pollen tubes germinate from the pollen grain and elongate through the pistil to deliver sperm cells to the egg cell. Once the pollen tube reaches the ovule, it discharges sperm cells to fertilize the egg cell within the ovule [5][6][7]. Changes in Ca^2+^ levels occur in both male and female tissues during the reproductive stage. In females, Ca^2+^ concentrations within the pistil fluctuate during developmental stages, reaching their peak at the stage of anther dehiscence [8]. Moreover, in petunia, pollination triggers an increase in Ca^2+^ concentration within the pistil [9]. In males, the Ca^2+^ concentration forms a gradient and oscillates with a period of several tens of seconds at the apex of the pollen tubes, synchronized with their growth rate [10] [11]. The Ca^2+^ gradient is established by Ca^2+^ channels, including CNGC18 [12] and GLR1.2/ 3.7 [13], which are localized in the apical region of pollen tubes. The MILDEW RESISTANCE LOCUS O (MLO) family proteins, including MLO1, 5, 9, and 15, also function as Ca^2+^ influx channels operating downstream of RALF4/19 signaling [14]. Ca^2+^ regulates membrane traffic within pollen tubes, including both endocytosis and exocytosis [15]. Fluctuations in Ca^2+^ concentration also regulate the polymerization and depolymerization of actin filaments through the ROP1-mediated RIC3/RIC4 pathway [16]. Thus, the regulation of Ca^2+^ concentration within the pollen tube is a crucial factor and is thought to be influenced by the surrounding environment during its elongation within the pistil. However, the impact of extracellular Ca^2+^ concentration on pollen tube elongation and guidance remains largely unknown. In lily, modulation of extracellular Ca^2+^ concentration has been reported to affect both pollen tube elongation and its oscillatory dynamics. The optimal Ca^2+^ concentration range in lily (0.1–1.0 mM) differs substantially from that in *Arabidopsis* [17].

Using live imaging of the Ca^2+^-sensitive fluorescent protein GCaMP6s[18], we examined whether changes in extracellular Ca^2+^ levels influenced the internal Ca^2+^ environment in the pollen tubes. We demonstrated that the extracellular Ca^2+^ concentration influences pollen tube morphology, growth rate, and guidance efficiency. We showed that high extracellular Ca^2+^ concentrations lead to elevated intracellular Ca^2+^ levels in pollen tubes, resulting in the disruption of the directional control of growth. Although the Ca^2+^ dynamics of the non-apical region of pollen tubes have been poorly characterized, the use of GCaMP6s has enabled their visualization. Oscillatory Ca^2+^ waves initiated at the apex of the pollen tube propagated toward the basal region, reaching the vicinity of the vegetative nucleus. Furthermore, pollen tubes that reached the ovule exhibited a pronounced increase in intracellular Ca^2+^ concentration. This was accompanied by an increase in the fluorescence intensity of Ca^2+^ waves. Our observations showed that external Ca^2+^ levels directly modulated the intracellular Ca^2+^ concentration within the pollen tubes.

## Methods

### Plant materials

*Arabidopsis thaliana Col-0* was used as the wild type. Plants were grown under 16hr light/8hr dark cycle at 23 °C. Pollen germination medium (PGM) was prepared as previously described [19]. Different CaCl_2_ final concentrations were tested as 1, 5 and 10 mM within the PGM.

### Plasmid construction and generation of transgenic plants

*RPS5Apro::H2B-tdTomato*/pMDC99 (coded as DKv039) was constructed previously[20]. *RPS5Apro::GCaMP6s*/pPZP211 (coded as DKv835) was constructed by replacing the 35S promoter of *35Spro::GCaMP6s*/pPZP211 (coded as DKv823)[21]. The binary vectors were introduced into *Agrobacterium* tumefaciens strain EHA105. The floral dip method was used for *Arabidopsis* transformation[22].

### Quantification of pollen tube growth rate and guidance rate

The pollen tube growth rate was measured using a semi-*in vivo* assay. [23]. The pollen tube growth rate was quantified under conditions without ovules. Cut styles were placed vertically on a cellulose cellophane layered on PGM agarose.

Observations were conducted using an inverted microscope (IX-71; Olympus) with a 10× objective lens (UPlanSApo, NA = 0.40; Olympus) and a CCD camera (DP73; Olympus). DIC images were taken every 10 minutes. The growth rate was quantified using the MTrackJ plug-in [24] in the Fiji software[25]. To quantify the guidance rate, cut styles were positioned horizontally on PGM agarose, and 6–7 ovules were arranged along the direction of pollen tube growth, extending from each style. The guidance rate was quantified as previously described [19].

### Kymograph

Kymographs of GCaMP6s and H2B-tdTomato fluorescence were drawn using the KymographBuilder plug-in (https://imagej.net/plugins/kymograph-builder) in Fiji software. Ca^2+^ concentration was visualized as a heat map, with the nucleus shown in magenta.

### Imaging pollen tube Ca^2+^ dynamics

Calcium dynamics within pollen tubes were observed using an inverted microscope (IX-81; Olympus) equipped with a spinning disk confocal unit (CSU-X1; Yokogawa Electric), 488 nm and 561 nm LD lasers (Sapphire; Coherent), and an EM-CCD camera (Evolve 512; Photometrics) using 20× (UPLFLN 20X, NA = 0.50; Olympus) and 60× objective lenses (UPLSAPO60XS, NA = 1.30; Olympus). We used two band-pass filters: 520/35 nm for GCaMP6s and 600/37 nm for tdTomato. The images were processed using Metamorph (Universal Imaging). Images were taken every 30 seconds for Fig. 2, every 3 seconds for Fig. 3, and every 10 seconds for Fig. 4.

**Fig. 1.**
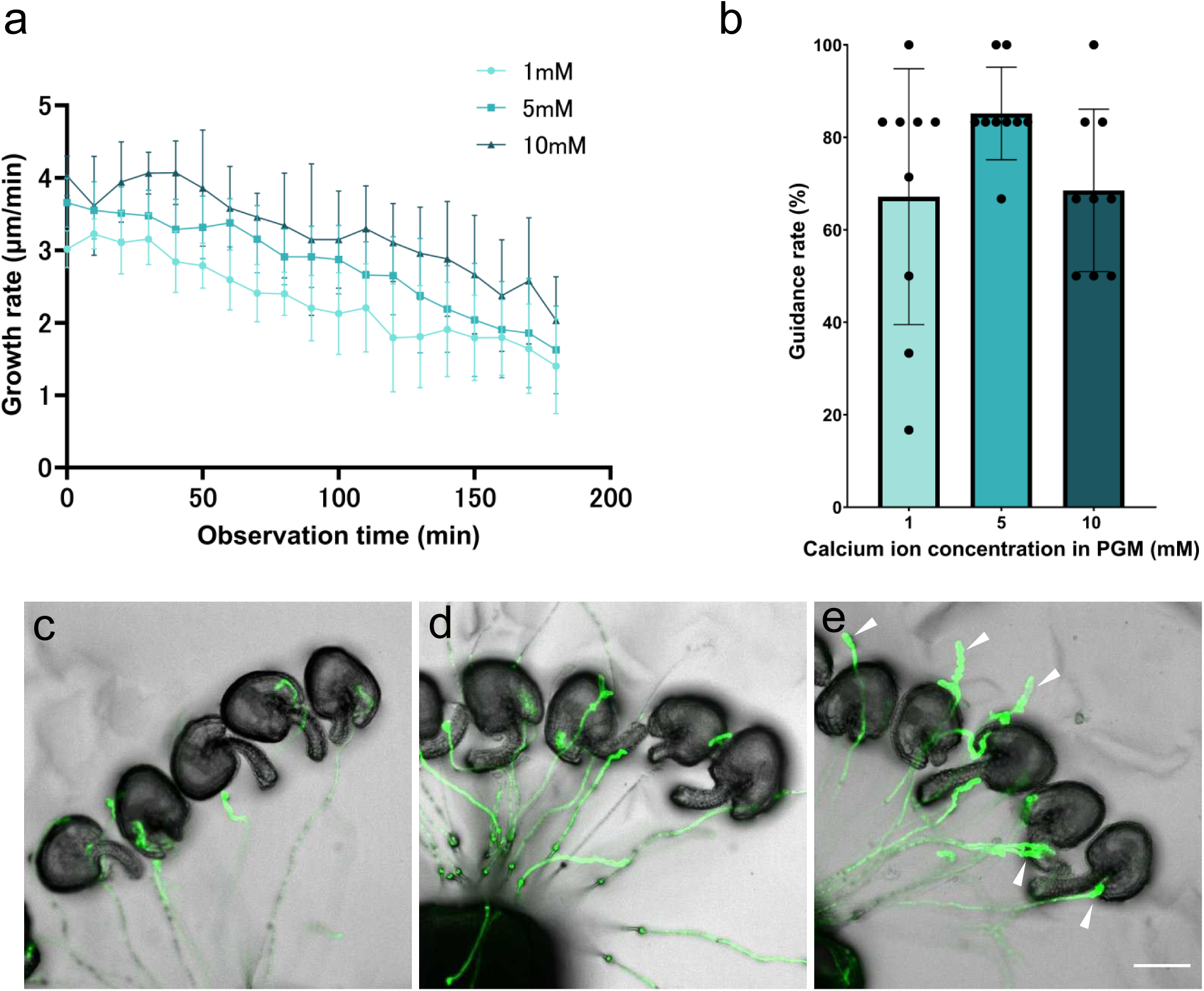
Effects of Ca^2+^ concentration in pollen germination medium (PGM) on pollen germination. (a) Quantification of pollen tube growth rate. Wild-type styles were pollinated and placed vertically on PGM agarose. Images were taken every 10 minutes under 1, 5 and 10 mM Ca^2+^. Data are shown as mean ± standard deviation (SD). [n= 22 (1 mM Ca^2+^), n= 24 (5 mM Ca^2+^) and n= 17 (10 mM Ca^2+^)]. (b) Quantification of pollen tube guidance rate in semi-*in vivo* assay. The percentage of ovules targeted by pollen tubes expressing GFP in the cytosol was measured (6-7 ovules per style). The total number of ovules was 55 (1 mM Ca^2+^), 54 (5 mM Ca^2+^), and 54 (10 mM Ca^2+^). Data are presented as mean ± standard deviation (SD). (c) –(e) Typical example of pollen tube guidance assay with 1 mM (c), 5 mM (d), and 10 mM Ca^2+^ (e). Arrowheads in (e) indicate coiling pollen tubes. Scale bar; 100 μm.

**Fig. 2.**
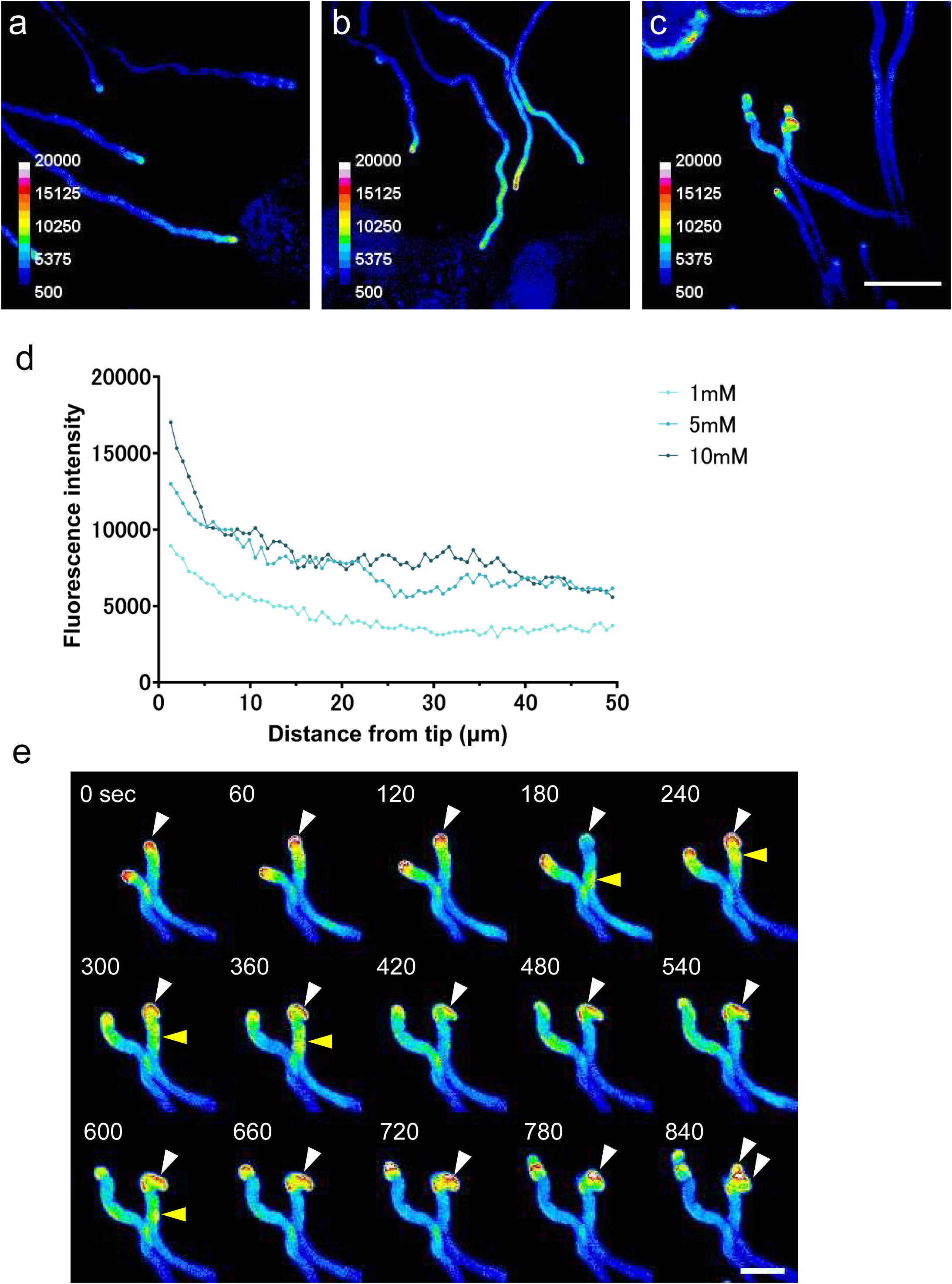
Ca^2+^ concentration in the medium and fluorescence intensity of the Ca^2+^-sensitive fluorescent protein GCaMP6s within the pollen tube. Images of pollen tube guidance assay with 1 mM (a), 5 mM (b), and 10 mM Ca^2+^ (c). Scale bar;100 μm. (d) Measurement of GCaMP6s fluorescence intensity of pollen tube tip region under different Ca^2+^ concentrations. The average fluorescence intensities from the tip to 50 μm were measured (n= 3 for each sample). (e) Time-lapse observation of pollen tubes expressing GCaMP6s in 10 mM Ca^2+^ medium. White arrowheads indicate the tip region with the maximum Ca^2+^ concentration. Yellow arrowheads indicate the Ca^2+^ waves in the shank region that propagated from the tip. The pollen tubes showed a coiled growth pattern near the ovules. Scale bar; 20 μm.

**Fig. 3.**
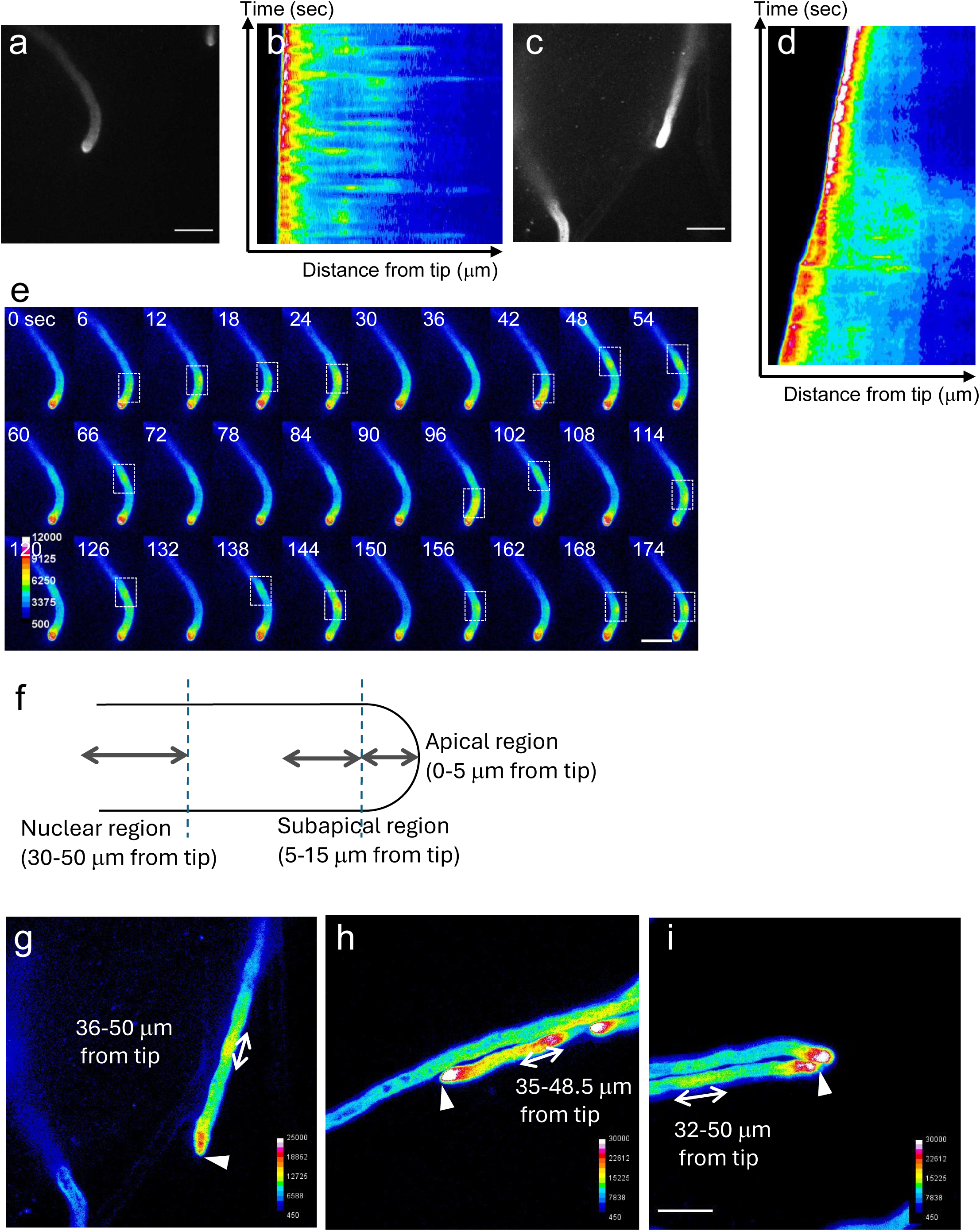
Ca^2+^ oscillations and wave propagation in pollen tubes. (a) Pollen tube growing toward the ovule. (b) Kymograph showing GCaMP6s fluorescence intensity along the pollen tube growing axis in (a). Horizontal axis, distance from tip (8.91 µm); vertical axis, time (195 sec). (c) Another example of a pollen tube growing toward the ovules. (d) Kymograph showing GCaMP6sfluorescencet intensity along the pollen tube growing axis in (c). Horizontal axis, distance from tip (17.79 µm); vertical axis, time (420 sec). (e) Time-lapse images are shown as montages. The white dashed boxes show the Ca^2+^ wave propagation from the tip. (f) Zonation of the pollen tubes was defined as follows in this study: apical region (0-5 μm from the tip), subapical region (5-15 μm from the tip), and nuclear region (30-50 μm from the tip). (g) – (i) Ca^2+^ wave propagation from the apex to the nuclear region. White arrowheads indicate the positions of the pollen tube tips. Scale bars; 20 μm.

**Fig. 4.**
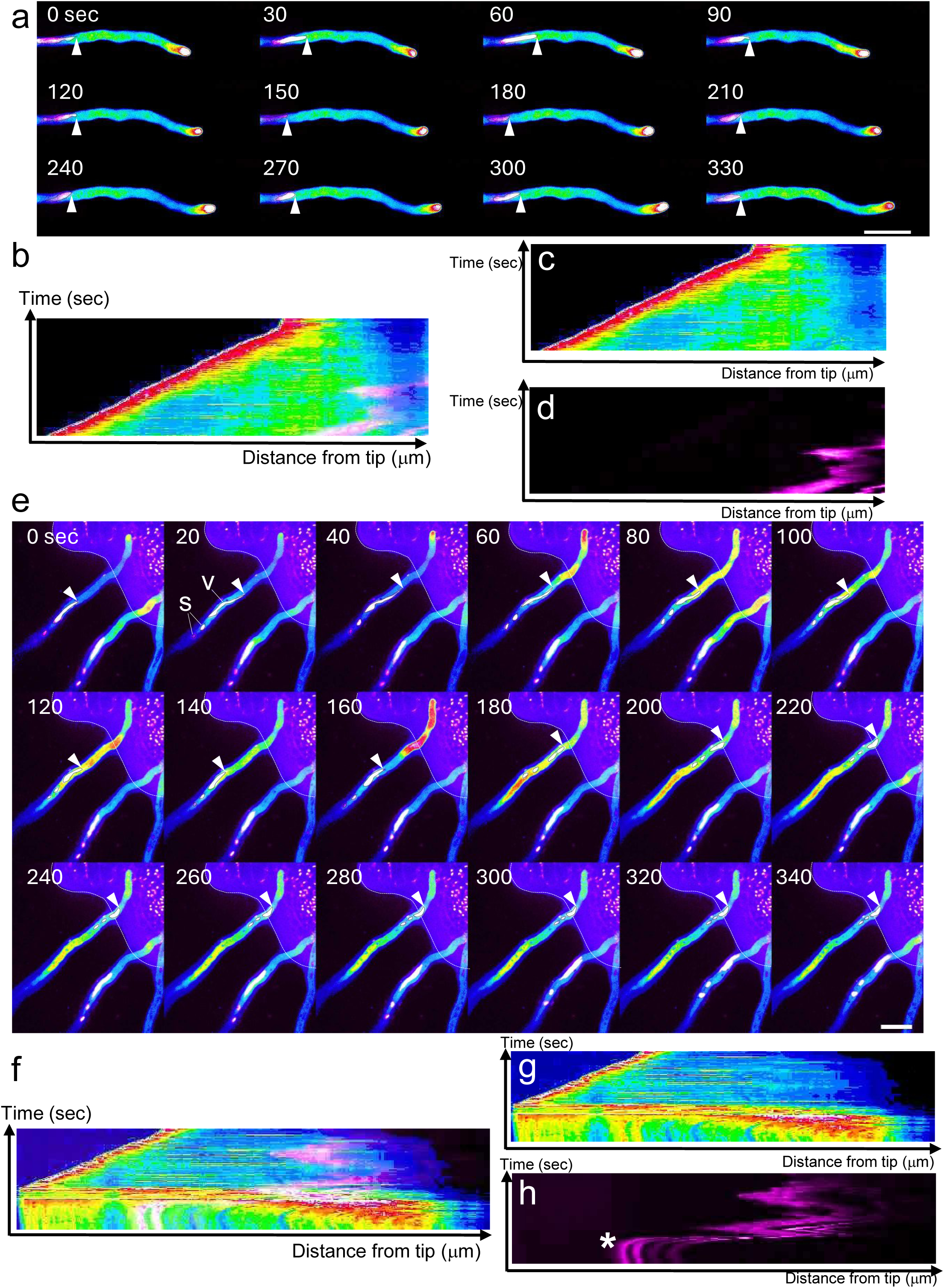
Ca^2+^ wave propagation and nuclear movement in pollen tube. (a) Pollen tube expressing GCaMP6s (heatmap) and H2B-tdTomato (magenta). Ca^2+^ wave reached approximately the tip of the vegetative nucleus. The vegetative nucleus is delineated by black dashed lines. White arrowheads indicate the leading edges of vegetative nuclei. (b) Kymograph showing GCaMP6s (heatmap) and H2B-tdTomato (magenta) fluorescence intensity along the pollen tube growing axis in (a). Horizontal axis, distance from tip (94.16 µm); vertical axis, time (363 sec). (c) Kymograph showing GCaMP6s (heatmap) fluorescence intensity along the pollen tube growing axis in (a). (d) Kymograph showing H2B-tdTomato (magenta) fluorescence intensity along the pollen tube growing axis in (a). (e) Pollen tubes expressing GCaMP6s (heatmap) and H2B-tdTomato (magenta) reaching the ovules. Ca^2+^ waves showed higher intensity than those in pollen tubes growing toward the ovules. The vegetative nucleus (v) and sperm cells (s) are delineated by black dashed lines. The white arrowheads indicate the leading edge of the vegetative nucleus. The vegetative nucleus and sperm cells move forward with strong Ca^2+^ wave propagation. White dashed lines delineate the ovule outline. (f) Kymograph showing GCaMP6s (heatmap) and H2B-tdTomato (magenta) fluorescence intensity along the pollen tube growing axis in (e,). Horizontal axis, distance from tip (110.88 µm); vertical axis, time (306 sec). (g) A kymograph showing GCaMP6s (heatmap) fluorescent intensity along pollen tube growing axis in (e). (h) A kymograph showing H2B-tdTomato (magenta) fluorescent intensity along pollen tube growing axis in (e). An asterisk indicates the point at which rapid nuclear migration occurred. Scale bars; 20 μm.

## Results

### Pollen tube growth was influenced by external environmental Ca^2+^ conditions

The intracellular Ca^2+^ concentration in the pollen tube is regulated by calcium channels at the apical region[12][13] [14][26]. While Ca^2+^ and reactive oxygen species (ROS) are thought to be supplied from the pistil during pollen tube elongation, the influence of the external Ca^2+^ environment on this process remains unexplored in the model plant *Arabidopsis thaliana*. To this end, we examined the effects of varying Ca^2+^ concentrations in the growth medium on pollen tube elongation and guidance. Following pollination of the pistil, the style was excised above the ovary, and the growth rate of pollen tubes emerging from the style was quantitatively examined in relation to the Ca^2+^ concentration of the growth medium. Elongation was faster at higher Ca^2+^ concentrations at any time during the observation period (two to five hours after pollination), as demonstrated by comparisons among 1-, 5-, and 10-mM treatments (Fig. 1a). These results indicate that external Ca^2+^ uptake plays a critical role in regulating the pollen tube growth rate.

### The Ca^2+^ concentration in the growth medium affected pollen tube guidance and growth dynamics

Subsequently, we investigated the effect of extracellular Ca^2+^ concentration on the guidance of pollen tubes toward the ovules. Using a semi-*in vivo* (SIV) assay with ovules, we quantitatively assessed the proportion of ovules successfully targeted by pollen tubes. For each pistil, six to seven ovules were arranged to assess the pollen tube targeting efficiency. The highest pollen tube guidance efficiency was observed at a Ca^2+^ concentration of 5 mM (Fig. 1b). Furthermore, the Ca^2+^ concentration modulated the dynamics of pollen tube elongation during guidance near the ovule (Fig. 1c-e). While pollen tubes were efficiently guided at 5 mM Ca2 +, they exhibited pronounced three-dimensional coiling after reaching or passing through the ovules under 10 mM Ca2 + conditions (Fig. 1e, arrowheads). These findings suggest that under high Ca^2+^ conditions, pollen tubes may lose proper control over their growth direction following the perception of attractant cues.

### The Ca^2+^ concentration in the growth medium influenced Ca^2+^ accumulation at the apical region of the pollen tubes

To examine the effect of Ca^2+^ concentration in the medium on pollen tubes, we employed GCaMP6s, a Ca^2+^-sensitive fluorescent reporter protein characterized by high sensitivity and slow decay kinetics (Fig. 2). GCaMP6s was expressed in pollen tubes under the control of the *RPS5A* promoter [27]. Consistent with the findings in Fig. 1, variations in Ca^2+^ concentration in the medium led to distinct patterns of pollen tube elongation near the ovules. Pollen tubes exhibited relatively straight growth at 1 mM Ca^2+^, displayed undulating trajectories at 5 mM Ca^2+^, and showed pronounced coiling at 10 mM Ca^2+^ (Fig. 2a-c). Quantitative analysis revealed that the GCaMP6s fluorescence intensity at the pollen tube apex increased with higher Ca^2+^ concentrations in the growth medium. Notably, the most significant differences in fluorescence intensity were observed at the extreme apex of the pollen tube (Fig. 2d). Time-lapse imaging of pollen tube elongation near the ovule in 10 mM Ca^2+^ medium revealed a repeated pattern in which a transient increase in Ca^2+^ concentration at the tip was followed by lateral Ca^2+^ accumulation, directing growth toward one side and ultimately leading to coiling (Fig. 2e, white arrowheads, and Supplementary Mov. S1). Moreover, the propagation of tip-localized GCaMP6s fluorescence signals toward the basal region of the pollen tube was also observed (Fig. 2e, yellow arrowheads).

### Ca²⁺ oscillations propagated from the apex to the nuclear region

The Ca^2+^ concentration at the pollen tube tip exhibits oscillations with a period of several tens of seconds, accompanied by corresponding oscillations in the pollen tube growth rate[11][28]. Prompted by the observation in Fig. 2e that tip-localized fluorescence propagated toward the base, we conducted live imaging analysis to investigate the Ca^2+^ dynamics. Compared with GCaMP6f [29], GCaMP6s offers the advantage of detecting weaker Ca^2+^ signals. Although it exhibits slower decay kinetics, it remains suitable for delineating active regions, similar to GCaMP6f. The temporal changes in the fluorescence intensity of GCaMP6s in the apical regions shown in Fig. 3a and 3c were quantitatively measured and displayed as kymographs (Fig. 3b and d, respectively).Consistent with prior studies [11][30][31], oscillatory Ca^2+^ dynamics were detected at the apex of the pollen tube (Fig. 3b and 3d). Furthermore, we identified a novel phenomenon in which wave-like Ca^2+^ signals propagated toward the basal region (Fig. 3a-d). In Fig. 3e, the white dashed box highlights the peak of Ca^2+^ waves. Following the peak of GCaMP6s fluorescence oscillation at the tip, Ca^2+^ waves propagated toward the basal region with a temporal delay (Supplementary Mov. S2). Wave-like Ca^2+^ fluorescence was observed in the nuclear region, located 30–50 μm behind the apex (Fig. 3g-i). Earlier studies have described this phenomenon as a “flicker” rather than a wave, likely reflecting the differences in the properties of GCaMP6s and GCaMP6f [32]. Given that the GCaMP6s used in this study is more sensitive to weak Ca^2+^ signals and shows slower decay of fluorescence than GCaMP6f, it is likely that the observed oscillatory Ca^2+^ dynamics were not restricted to the tip region alone.

To assess Ca^2+^ dynamics in the nuclear region, we performed dual labeling of the nucleus and Ca^2+^ signal. To visualize the nucleus, we employed a construct expressing H2B-tdTomato under the control of a ubiquitously expressed *RPS5A* promoter. The Ca^2+^ wave reached the area near the leading edge of the vegetative nucleus (shown in magenta in Fig. 4a-d and Supplementary Mov. S3). As illustrated in the kymograph in Fig. 4d, the nucleus exhibited repeated forward and backward movements within the pollen tube [33]. The Ca^2+^ wave propagates to the vicinity of the vegetative nucleus but does not extend further toward the basal region, possibly due to physical hindrance by the nucleus.

### Ca^2+^ concentration within the pollen tube increases as it approaches the ovule

As previously reported [34], the overall Ca^2+^ concentration in the pollen tube increased as it approached the ovule (Fig. 4e-h). Although Ca^2+^ waves typically exhibit weaker fluorescence than the apex, the fluorescence levels in the pollen tubes near the ovule were comparable to those at the tip. As the pollen tube reached the ovule and Ca^2+^ levels surged, the nucleus migrated toward the apex of the pollen tube. At 160-180 sec, Ca^2+^ burst within pollen tubes caused rapid migration of the vegetative nucleus and sperm cells toward the apical region (Fig. 4e, arrow heads; Fig. 4h, asterisk). These findings suggest a correlation between Ca^2+^ dynamics within the pollen tube and nuclear migration.

## Discussion

During pollen tube elongation through the pistil, the Ca^2+^ environment within the transmitting tracts is predicted to exert a significant influence on both growth and guidance. However, no study has visualized this effect using live imaging approaches. This study employed a semi-*in vivo* guidance assay to enable live imaging of Ca^2+^ dynamics under conditions that closely mimic the *in vivo* environment. Ca^2+^ imaging was performed using GCaMP6s, a highly responsive Ca^2+^ -sensitive fluorescent protein. As expected, the Ca^2+^ concentration within the pollen tube reflected the Ca^2+^ concentration in the external environment. In *Arabidopsis*, pollen tube elongation accelerates with increasing Ca^2+^ concentrations up to 10 mM, whereas in lily, optimal elongation is observed at Ca^2+^ concentrations ranging from 0.1 to 1 mM [17]. The differing growth environments, through a hollow pistil in lily versus intercellular spaces in *Arabidopsis*, are likely to result in distinct physiological requirements. With respect to guidance, high Ca^2+^ concentrations disrupt the directional control of pollen tube elongation. The Ca^2+^ wave identified for the first time in this study had previously appeared only as transient “flickers” in experiments employing the faster-response indicator, GCaMP6f [32]. Using GCaMP6s, which exhibits slower signal decay, we were able to track and detect weak Ca^2+^ signals over time. Ca^2+^ waves are also observed in neurons, where they play a role in regulating the extension of axons and dendrites [35][36]. In neurons, Ca^2+^ waves can propagate over long distances through the release of Ca^2+^ from intracellular stores such as the endoplasmic reticulum [37]. The mechanism by which Ca^2+^ waves propagate in pollen tubes remains to be elucidated and requires further investigation.

When extracellular Ca^2+^ levels are elevated, an excessive influx of Ca^2+^ into the cell occurs through calcium channels. In the presence of 10 mM Ca^2+^, pollen tubes displayed abnormal morphology, with three-dimensional coiling. These observations suggest that excessively high Ca^2+^ concentrations at the pollen tube tip disrupt polar growth, causing lateral extension toward regions with relatively lower Ca^2+^ concentrations, after which directional growth becomes increasingly unstable. Because coiling occurred irrespective of ovule presence, it is likely attributed not to attractant signals from female tissues but to elevated intracellular Ca^2+^ levels in the pollen tube.

The Ca^2+^ level reached its maximum at the tip of the pollen tube and subsequently propagated basally, arriving at the leading edge of the vegetative nucleus. The reorganization of the actin network triggered by the Ca^2+^ wave may be essential for nuclear migration associated with pollen tube elongation [11]. Although the Ca^2+^ wave was not observed beyond the vegetative nucleus toward the basal region, it is possible that the vegetative nucleus functions as a physical barrier.

Upon reaching the ovule, pollen tubes exhibit a sharp increase in Ca^2+^ concentration, which, once propagated near the nuclear region, triggers a rapid relocation of the vegetative nucleus and sperm cells toward the tip. A future objective is to determine whether nuclear movement or release is related to Ca^2+^ signaling. This question could be elucidated by imaging Ca^2+^ dynamics and sperm cell release in the pollen tubes of mutants with restricted nuclear movement.

### Concluding remarks

The external environment of the pollen tube affects intracellular Ca^2+^ levels, as well as pollen tube growth and guidance, suggesting that communication between the pollen tube and pistil plays a crucial role in successful reproduction. These findings further imply that Ca^2+^ dynamics may contribute to the regulation of nuclear migration in pollen tubes.

## Contribution statement

K.M.T. designed the study and performed the experiments. Y.M. and D.K. designed and prepared the plasmid constructs. T.H. provided advice on the experimental design and data interpretation. All authors discussed the results and wrote the manuscript.

## Acknowledgements

We would like to thank Yoshikatsu Sato (Institute of Transformative Bio-Molecules, Nagoya University) for support in utilizing microscopes. This manuscript was prepared with the assistance of ChatGPT and Paperpal to improve language editing and clarity. The authors have reviewed the manuscript and are fully responsible for the scientific content.

